# Template-based design of peptides to inhibit SARS-CoV-2 RNA-dependent RNA polymerase complexation

**DOI:** 10.1101/2022.01.24.477502

**Authors:** Akshay Chenna, Wajihul H Khan, Rozaleen Dash, Anurag S Rathore, Gaurav Goel

## Abstract

The RNA-dependent RNA polymerase (RdRp) complex of SARS-CoV-2 lies at the core of its replication and transcription processes. The interfaces between the subunits of the RdRp complex are highly conserved, facilitating the design of inhibitors with high affinity for the interaction hotspots of the complex. Here, we report development and application of a structural bioinformatics protocol to design peptides that can inhibit RdRp complex formation by targeting the interface of its core subunit nonstructural protein (nsp) 12 with accesory factor nsp7. We adopt a top-down approach for protein design by using interaction hotspots of the nsp7-nsp12 complex obtained from a long molecular dynamics trajectory as template. A large library of peptide sequences constructed from multiple hotspot motifs of nsp12 is screened *in silico* to determine peptide sequences with highest shape and interaction complementarity for the nsp7-nsp12 interface. Two lead designed peptide are extensively characterized using orthogonal bioanalytical methods to determine their suitability for inhibition of RdRp complexation and anti-viral activity. Their binding affinity to nsp7 (target), as determined from surface plasmon resonance (SPR) assay, is found to be comparable to that of the nsp7-nsp12 complex. Further, one of the designed peptides gives 46 % inhibition of nsp7-nsp12 complex at 10:1 peptide:nsp7 molar concentration (from ELISA assay). Further optimization of cell penetrability and target affinity of these designed peptides is expected to provide lead candidates with high anti-viral activity against SARS-CoV-2.

## Introduction

The Coronavirus disease 2019 (COVID-19) is caused by a new strain of *β*-coronaviruses termed Severe Acute Respiratory Syndrome Coronavirus 2 (SARS-CoV-2).(1, 2) At the heart of the transcription machinery of SARS-CoV-2 virus is the RNA-dependent RNA polymerase (RdRp) which controls the genomic replication processes of single stranded RNA viruses. The genome is used as a template by hijacking the machinery of the host cells to translate RdRp which in turn is used to complete the transcriptional synthesis of different protein structures and RNAs in SARS CoV2.(3, 4) RdRp is a trimeric complex of three different proteins non structural proteins viz NSP7, NSP8 and NSP12 of which NSP12 is the core catalytic unit and a target of several drug discovery programs.(5, 6) Recent studies revealed highly conserved structural and functional features of RdRp in coronaviruses and an amino acid sequence identity of 96% with the RdRp of SARS-CoV.(7, 8) It is understood that the interaction of NSP7 and NSP8 with NSP12 significantly enhances the polymerase activity of the otherwise minimal activity of innate NSP12. (9–11) Therefore targeting the integrity of the RdRp complex by exploring the hotspots to disrupt the protein-protein interactions in the subunit has been suggested as an effective drug discovery strategy.(12, 13) The crystal resolved structure and long trajectory from Molecular Dynamics (MD) simulations of RdRp reveal major contributions to NSP12 binding by NSP7 as opposed to NSP8 which forms much fewer contacts.(6, 14) Cryo-EM maps revealed that the N-terminal region of NSP8 adopts an extended, disordered conformation making it a challenging target for disrupting protein-protein interactions (PPIs). Moreover the binding site on NSP12 made by NSP7 is well conserved in contrast to the binding site by the NSP8 subunit.(7) NSP7 in SARS-CoV-2 shares 100% sequence similarity with SARS-CoV in stark contrast to the envelope proteins of coronaviruses. Additionally, NSP7 is found to make several protein-protein interactions in the cellular viral proteome making it a prime pharmacological target.(15) Three FDA approved small molecules drugs viz. Metformin, Entacapone and Indomethacin were identified with the potential role of disrupting the network of protein interactions made by NSP7. (13) However, none of these drugs target the interface made with NSP12 protein. Therefore we have chosen the protein-protein interface of NSP12-NSP7 complex as a target for development of orthosteric inhibitory drugs.

Peptide sequences can be computationally tailored to mimic the hotspot interactions of one of the binding partners and are thus considered as natural inhibitors of PPIs.(16, 17) Several peptides have been reported as potent against microbial pathogens including the recent approval of anti-HIV peptides.(18–20) With a steady development and approval of peptide based drugs in the recent past, they are seen as promising alternatives to small molecule drugs due to high selectivity and easy of manufacturing.(21) Short peptides possess low immunogenicity profiles, minimal off-target interactions and cheaper production costs. Despite their effectiveness in disrupting PPIs, peptides have a notably short duration of action due to proteolysis and rapid renal clearance.(22) Protein based drugs such as peptides, miniproteins, nanobodies and antibodies have also been identified to target PPIs of structural proteins on SARS-CoV2.(21, 23–29) There are over 60 small molecules targeting the enzymatic activity of RdRp and over 30 small molecules targeting intracellular PPIs are in active clinical trials (30) yet fewer peptide based drugs have been developed for intracellular targets.(6, 31) In this work, we report short peptide sequences that bind with nanomolar affinity (~100 nM) to the NSP7 protein monomer of RdRp to inhibit the polymerase activity of NSP12. Protein-protein interaction hotspots identified from a long molecular dynamics (MD) trajectory were used as a template for creating an *in-silico* library of peptide sequences. Our in-house structural bioinformatics based protocols mined and iteratively screened peptide sequences to generate valid decoys that bind with similar or slighlty higher affinity than the original protein receptor.

## Results and discussion

### Determination of hotspot residues on NSP12 interface

Heteromeric protein-protein interactions bury on an average 1900 Å^2^ of surface area upon binding, translating to 57 amino acids per interface.(32) However only a small fraction of these residues at the protein interface contribute the largest to binding energy. These residues are termed as hotspots of binding interactions.(33) In our work we demonstrate a structural bioinformatics pipeline for designing peptides by identifying hotspots from a long molecular dynamics (MD) trajectory. Our design strategy begins with using the information from identified hotspots from one of the protein binding partners as templates for generation of a library of peptide sequences with a goal of antagonistically inhibit a protein-protein interaction. Similar template based strategies involve using the hotspot residues from the cystal complex of the heterodimer to construct a optimised protein,(16) or from the topologocial information of the binding partner without using the sequence.(24, 27) By aiming to mimick the binding interactions of NSP12 we seek to achieve equal or improved bindng affinity to the target protein (NSP7) at optimal concentrations to inhibit *in-vivo* viral interactions.(34) Peptide mimetics of protein protein interactions present several advantages. Firstly, it understood that peptides derived from the binding interface can mimic the entire partner protein and bind more tightly to the target protein.(35) An exhaustive study carried out using X ray crystallography and rigid-body ligand docking found out that the interface of the holo protein in the ligand bound state shows a high match with the interface in a protein-protein complex.(36) This study corroborates the motive for conformational change induced upon ligand binding to mimic that of the partner protein. The results from their work thus suggest that the protein-bound conformation of the receptor is a significantly better starting point for drug design than the *apo* structure. London et al. have demonstrated on a large scale that self-inhibitory peptides can be derived from the interfaces of protein-protein interactions.(37) These peptides have been shown to be effectively mimic the binding modes of the origin domain of the peptide and bind with similar or better binding affinities owing their origin to *hot segments* on the protein-protein interface.(38) Secondly, it has been shown that the conformational change upon binding to helical proteins results on an average of 0.11 nm change in the RMSD of the C*α* atoms compared to the free state, resembling the *apo* conformation in the *holo* state.(39) Since NSP7 is an helical bundle of three helices, we believe that peptides derived form the bound interface of NSP12 will effectively bind to the *apo* state of NSP7. Additionally, computational solvent mapping studies which estimate the ligand druggability demonstrate that the interactions between globular proteins results in conformational changes largely restricted to 0.6 nm of the binding hotspot representing a high degree of structural conversation of the binding hotspot in the *apo* and *holo* states.(40) Another study demonstrates that surfaces with binding sites are predisposed in the *apo* structure of globular proteins making them a feature of druggable sites as found from computational fluctuational simulations.(41) Thirdly, from a more realistic statistical ensemble picture it can be argued that the fit induced in NSP7 upon binding to NSP12 results in a conformation that is already present in the apo state of NSP7 since binding only induces a shift in the relative population of the conformations that favour binding.(42) This justifies our use of the *holo* state for designing effective inhibitors of PPIs. We determined the hotspot residues on NSP12 in the NSP12-NSP7 dimer subunit complex from a MD trajectory of RNA-dependent RNA polymerase (RdRp) of SARS-CoV2 by calculating the difference in the ressidue-wise area buried (eq 1). The trajectory was clustered using a cutoff of 2.5 Å on the backbone atoms of Rdrp. The residue-wise hotspot areas were weighted (Δ*A*_*i*_) by the population fraction of the cluster.

As a result, the interface residues on NSP12 contribute differently to binding with NSP7 indicating a complex, discontinuous binding epitope (Figure 1). The interface was split into contiguous stretches of amino acid sequences by allowing upto three intervening non-interface residues (i.e. Δ*A*_*i*_ = 0). A similar approach was followed by Jones and Thornton (43) allowing a break of five residues. Paletal. (44) analyzed interfaces of heterodimeric complexes and found that a typical surface contains an average of 5.6 segments in agreegment with our observation of five segments on the interface o f NSP12 of which the prominent three are highlighted in Figure 1. However allowing a shorter stretch of non-interface residues resulted in large segments consisting upto 20 amino acids.(SI)

**Fig. 1.**
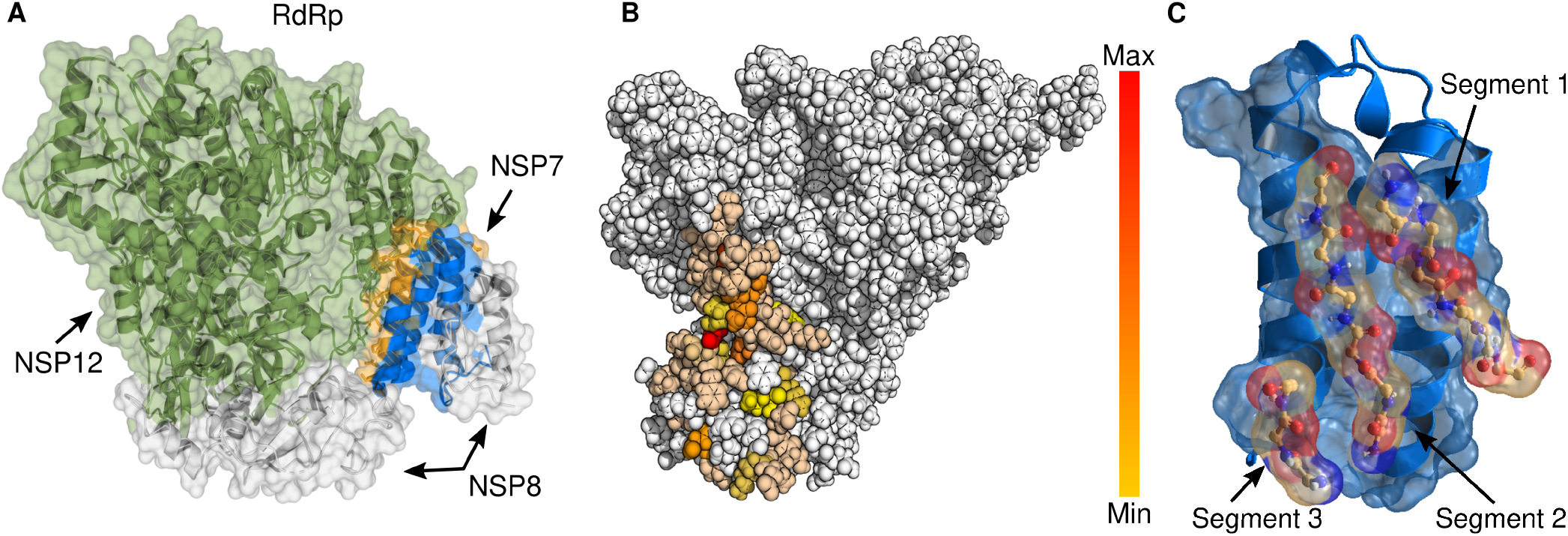
NSP12-NSP7 subunit interactions of RNA dependent RNA polymerase (RdRp). **A** Interactions between the proteins involved in RdRp complex are depicted. NSP7 is known to contribute the maximum to the binding affinity with NSP12. **B** Key hotspot interactions made by NSP12 as an ensemble average over a MD trajectory are highlighted with a color gradient. **C** Three sequentially contiguous hot-segments identified from the hotspots are shown. These segments are spatially connected by linkers to generate *in-silico* peptide libraries.

### Peptide design

Most studies have focused on design from a linear template of binding motifs from one segment. The disembodied segments identified f rom the h otspot r egion of NSP12’s interface originating from the dynamic binding interface were connected by linkers to create a composite peptide topology (Figure 1. Amino acid fragments within a segment were assembled by varying the sequence lengths leads to multiple sequences. Glycine and alanine linkers combinatorially connect every possible spatially close residues from two different segments based on the C*α* distances of the joining residues (Table 1). We also generated sequences from the same segment that resulted in a library of over 93,000 sequences ranging from three to 37 amino acids. Sequences originating from the parent segments contain rim residues or show little contribution to the binding hotspots were removed by scoring the library peptides. As shown in Figure 2 the constructed sequences are scored by the sum of the hotspot areas of the residues as found in the origin domain and weighted by the cluster population from which the interface was analysed (Σ_i_A_i_P^c^, Section). We found that 21 of the highest scored 50 sequences have a length of 8-12 amino acids, an ideal size for therapeutic peptides. Design of longer peptides is an added challenge as the folded states must be made stable enough to minimize conformational entropic cost upon binding to the target protein while very short peptides may not capture the native hotspot interactions made by NSP12.(39) The sequences derived are from the most populated cluster of the MD trajectory and contain no sequences from two segments.

**Table 1.** Residues from different segments were connected based on the C*α*-C*α* distances by a Glycine/Alanine bridge.

| Distance (nm) | Bridge |
| --- | --- |
| <0.7 | -G- |
| 0.7-1.0 | -G-A- |
| 1.0-1.3 | -G-A-A- |

**Fig. 2.**
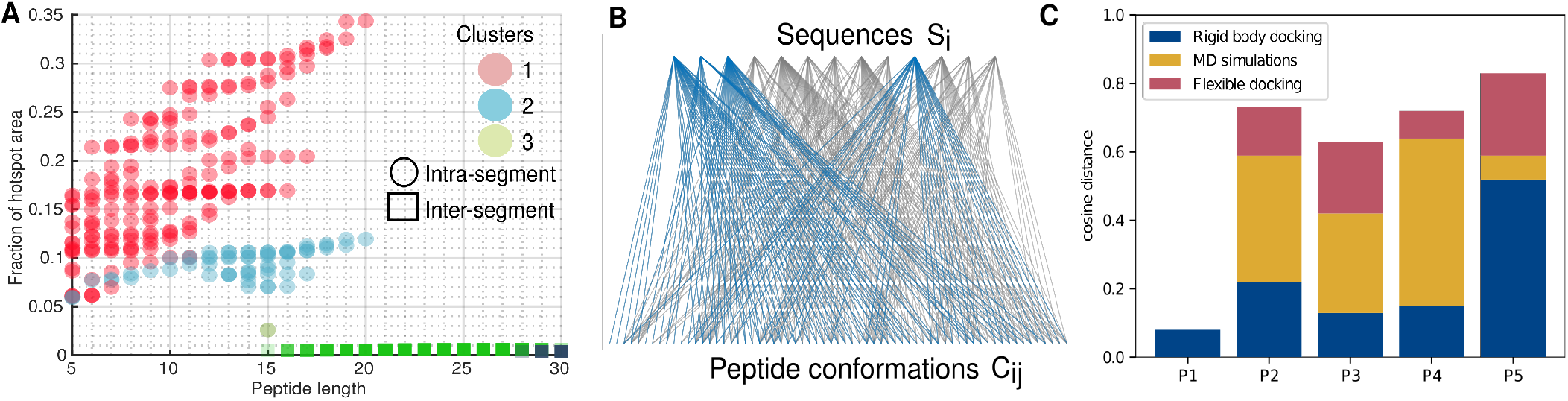
Bioinformatics and simulation strategies for screening peptides. **A** The library of peptide sequences constructed are scored with the hotspot areas of NSP12 segments identified from simulations. The plot compares the cumulative hotspot area of the peptide as a function of peptide lengths. Peptides derived from the same segment are shaped in º and inter-segment peptides are depicted in ◻. The sequences are colored with respect to the NSP12-NSP7 cluster from which the sequences were constructed. **B** TM-scores are cross computed for 65 peptide conformations generated from 13 peptide sequences are shown in the bottom and top layers respectively. An edge represents the ability of the peptides to adopt a conformation. Unique peptide sequences and folds identified from integer linear programming on a binary matrix of TM-scores are shown in blue. **C** Binding interface similarity measured by the cosine distance of the docked peptides in comparison to the interface of cluster center of the most populated cluster of NSP12-NSP7 is plotted for peptides (P1 to **P5**). Higher dot products indicate better ability of the peptides to replicate the binding modes of NSP12.

We therefore added two inter-segment sequences with the highest hotspot scores to the exisiting set of 21 for further design evaluations. Five low energy conformations for each of the 23 peptides were generated using a de-novo coarse grained optimized potential for efficient structure prediction (OPEP) forcefield at physiological pH.(45) The corresponding charge on the peptide was found at physiological *pH* from residue *pK*_*a*_ values using the propka web-server.(46) The peptide conformations were modelled by taking the structure of the receptor (NSP7) into account, generating poses of peptide-protein at the given binding patch. However we filtered 10 peptides due to their inability to mimic the backbone topology of the functional motifs from the origin domain (backbone RMSD > 0.35 nm). To achieve high binding activity, the peptides must fold into states that replicate the binding modes of the defined template(47). Additionally, this ensures that our peptides have a high complementary shape to the target as seen in the native hotspot interactions leading to high functionally accurate mimetics. Finally, our combinatorial approach of assembling amino acids might result in large number of entries with high sequence identity that could lead to the same therapeutic functionality. To remove redundant peptide sequences (and consequently folds) in the remaining set of 13 peptides, we used a TM-score (template modelling score) metric to classify similarity.(48) The peptides were aligned prior to TM-score evaluations by standard Needleman–Wunsch algorithm with an custom identity mutation matrix (see SI table for matrix). This method aligns the sequences based on their identities instead of similarities. We applied weighted integer linear programming (wILP) method on the 65 × 65 binary TM-score matrix of 13 peptides (each peptide has 5 conformations) to select a minimal set of non-redudant peptides with the highest hotspot fraction of the blueprint interface (Section). Optimisation of peptide similarity has resulted in five distinct peptide sequences (Figure 2). The selected peptides possess nearly 40% of the hotspot area as compared to the origin domain.

### Peptide binding

To evaluate the quality of our designs we adopted a multi-step docking process. We began with a global blind docking of each of the five shortlisted peptides (**P1**-**P5**) to allow for a complete 3 dimensional exploration of the target surface. This exercise will narrow down on the binding site locally favoured by the peptides providing information on the binding specificity of the designed molecules. A robust quantitative descriptor developed by Anivash et al. (17) to evaluate the binding of quality given by the cosine distance (cos *θ*, see methods for details on construction procedure) was determined for the docked peptide-NSP7 complexes. Higher values of cos *θ* indicate preference for the peptides to share the same interface with NSP7 as shared by NSP12 in the RdRp subunit. The binding energies of peptide-protein binding for evaluating cos *θ* were calculated by MMPBSA-WSAS (Molecular Mechanics/Poisson Boltzmann/ Surface Area-Weighted solvent accessible surface area)(49, 50) method to compute the enthalpy and vibrational entropy. We notice lower values of cos *θ* during the first step of as the peptides were rigidly docked. However, peptides are very flexible and are known to adapt to the unbound structure of the receptor protein. Therefore we have allowed the peptides to undergo local perturbations to allow the peptide to refold and optimise the binding. Initial hits (docked poses with highest cos *θ* with the exception of P1 rejected due to poor scores) were considered for this step of flexible docking. Using FlexPepDock protocol implemented in Rosetta we carried out small-scale rigid body motion of the peptide coupled with backbone shear.(51) Alternate binding modes were explored by enumerating mulitple side-chain rotamer conformations of individual residues of the peptide (Figure 2). By introducing flexibility we observe a significant improvement in the re-computed cos *θ*. This conformational sampling of the peptides is indicative of the ability of the peptides to fold and adapt a complementary shape to the backbone structure of the binding partner. As a final determinant of our design success we performed short all-atomic MD simulations of the peptide-protein complexes with the highest cos *θ* from the flexible docking stage in water buffer mimicking physiological conditions. Although we notice a decrease in the cosine similarity distance in the MD trajectories of the peptide-NSP7 complexes, the dot products can be considered to be high enough for the designed peptides to enable high specificity binding to the protein. Overall, considering the increasing intensity of validation with three docking steps we selected two designs viz. **P4** and **P5** for experimental validation as inhibitors of protein-protien association of the NSP12-NSP7 subunit of the RdRp complex.

### Experimental testing

RdRp functions as a polymerase in the infected cell’s cytoplasm. Thus drugs must penetrate the host cell to disrupt the intracellular PPIs of RdRp. Peptides **P2** to **P5** were appended by a short string of poly arginines at the N-terminal for evaluating cell-penetrability using sequence based predictors.(52) Sequences **P4** and **P5** were classified as probable cell-penetrable entites while sequences **P2** and **P3** showed poor cell-penetration confidence and consequently were not validated *in-vitro* for binding activity.(SI) We measured the kinetics of **P4** and **P5** binding to NSP7 via Surface Plasmon Resonance experiments (SPR, Figure 3B). Both peptides showed similar dissociation constants (K*D*) of 133 nM and 167 nM and similar binding kinetic profiles. The peptides possess marginally improved binding affinity with NSP7 in comparison to the NSP12-NSP7 complex (Table 3A). In contrast to the experimentally predicted binding energies, ΔG_MMPBSA-WSAS_ has over-estimated the the binding energies by nearly two orders of magnitude. Though it is noteworthy that the binding, free energies are predicted similar trends as measured by SPR. Peptide-NSP7 docked poses with the highest dot products (cos *θ*) after flexible docking are depicted in figure 3A with the interface patch of NSP7 with NSP12 is colored in yellow. Poses with high dot products signifying better binding specificity are a strong indicator of the peptides’ ability to mimic the binding modes of NSP12 and serve to validate our approach of peptide design. Subsequently, the peptides were tested for competitive binding with labelled NSP7 (his-taq) in the presence of immobilised NSP12 using ELISA binding assays. The signal in Figure 3B is a measure of the NSP7 bound to the NSP12. **P4** shows 46% inhibition in the binding of NSP7 with NSP12 at ten times the molar concentration of NSP7 translating to an IC _50_ of about 50 *μ*M for **P4**. However, **P5** did not show significant inhibition even at high concentrations indicative of specificity at a non-neutralising site. ELISA based competitive assays suggest that our peptide sequence needs further optimisation for achieving higher affinities, highlighting a fundamental shortcoming of our approach in restricting the sequences of peptides from the template. Similar observations were made by Valiente et al (27), where the L-isoforms showed a 43 times lower binding affinity to the spike protein in comparison to the D-isoforms. Moreover, by constraining the pepitdes to emulate the backbone fold of the template we are intrinsically limiting the maximum possible binding affinities of the peptides. (24, 34) *De-novo* design of stable folds could potentially enable topologies that manifest higher binding enthalpies and minimize entropic costs of folding to the holo state.

**Fig. 3.**
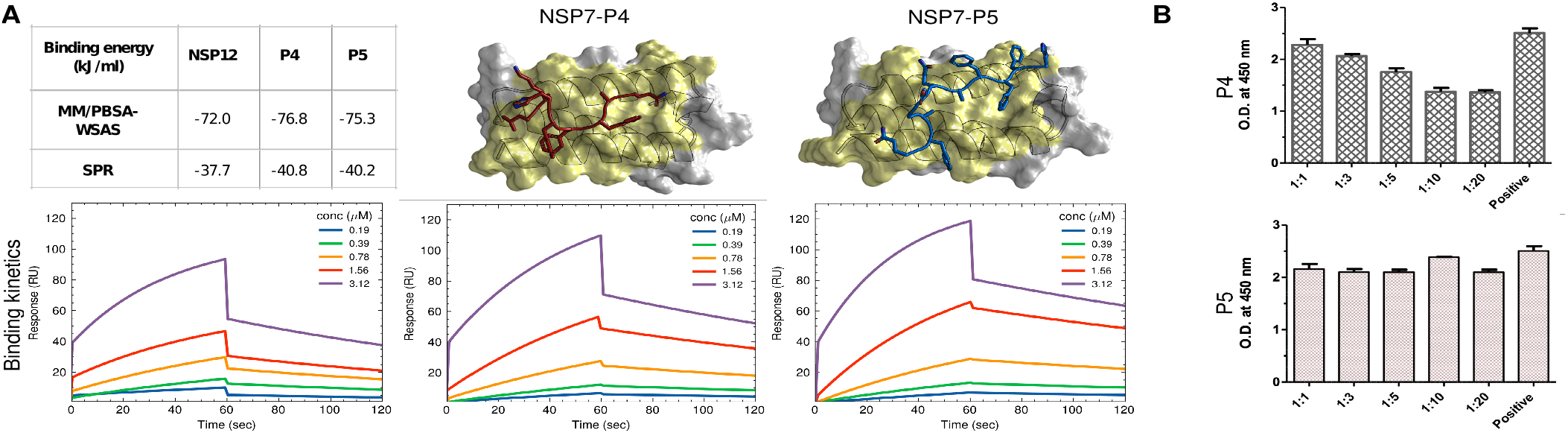
Peptide binding to NSP7. **A** Table compares the binding affinity of proteins from MMPBSA-WSAS method on the docked complex to the affinity determined from SPR experiments. SPR curves for NSP12-NSP7 and peptides **P4** and **P5** to NSP7 respectively. The cartoons show the binding pose of peptides to NSP7 with the highest cos *θ*. Patch colored in yellow is the interface of NSP7 when bound to NSP12. **B** Competitive inhibition of NSP12-NSP7 complexation in the presence of peptides is measured by ELISA assays. The signal is a measure of the NSP7 that remains bound to NSP12. Inhibition is measured for peptide ratios upto 20 times the concentration of NSP7.

## Conclusion

We developed a structural bioinformatics pipeline demonstrating rational design of peptide inhibitors of protein-protein interactions from a long MD trajectory. The protocols aim to identify valid sequences from a virtual library of sequences constructed using information of the template hotspots such that critical interactions made by the template are mimicked by the peptide. This is ensured by screening sequences containing a high fraction of the hotspot residues as identified in the template followed by their ability to reproduce the backbone folds from the structural repertoire of the interface of the template protein. To check the translation of the topological features of the peptides into binding affinity and specifity *in-silico*, the filtered sequences were docked in two stages, viz. rigid body global docking and flexible local docking. A final short MD simulation confirmed the presence of metastable peptide binding poses to the interacting interface of NSP7 in RdRp. Two sequences, **P4** and **P5** were shortlisted for experimental validation based on higher predicted cell-penetration confidence than **P2** and **P3**. Similar profiles of binding kinetics of the peptides and NSP12 to NSP7 from SPR measurements provide proof of concept of our approach in the ability of the peptides to replicate the origin domain interactions. Competitive assays based on ELISA substantiated the need for optimised sequences and folds different from the template for enhancing the binding affinity. Finally, we need to incorporate positive strategies for design that allows peptides to penetrate the cell membrane efficiently and target intracellular protein-protein interactions with high *in-vivo* efficacy. Requiring minimal computational efforts our template based design approach demonstrates that self inhibitory peptides derived from the interface of protein-protein interactions can serve as a good starting point for further refinement and lead optimisation.

## Methods and materials

### Computational design

#### Determination of binding hotspots

Key hotspot residue interactions made by NSP12 with NSP7 were obtained from the solvent accessible surface area analysis of its dimeric subunit complex of RdRp from a 10 *μ*s molecular dynamics trajectory. (14) Interface residues on NSP12 are found from Eq. 1

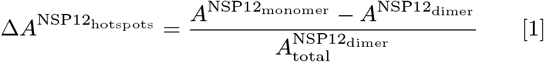

From a sufficiently long MD trajectory, a residue’s contribution to the binding hotspot is found by weighing its Δ*A* with a factor *Pc* that captures the probability of a residue to lie at the interface with NSP7. The MD trajectory was clustered using a 0.25 RMSD cutoff on the backbone atoms of the NSP12-NSP7 dimeric subunit. *Pc* for a given cluster *c* is the population fraction of the cluster 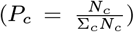 as observed in a MD trajectory obeying the ergodic principle.

#### Design strategy

Following the work of Mishra et al., (17) sequences of peptides were built by sewing a chain of spatially close hotspot residues from two different binding segments of the hotspot as given in Table 1. A segment is defined as a continuous sequence of residues on the NSP12 binding interface. The segment terminates if three or more consecutive residues are not part of the binding hotspot (i.e. *P*_*i*_Δ*A* = 0). A combinatorial library of such inter-segment connections was made and appended by intra-segment derived sequences. Peptides in the generated library were scored by the hotspot areas as evaluated from their parent NSP12 conformation. Peptide sequences with length less than 12 amino acids were selected from the top scoring sequences. Five conformations with the lowest energy were selected using the OPEP forcefield implemented in PEPFOLD. (53)

#### Assessment of peptide fold and elimination of redundancy

Peptides are rejected if the backbone of the lowest energy conformations do not adopt to their origin motif’s (NSP12) structure using a criteria of < 3.5 Å backbone RMSD. A structural template modelling score (TM-Score) was applied for evaluating conformational and sequence similarity in the shortlisted peptides.(54) The sequences of the selected set of peptides were aligned using the Needleman–Wunsch algorithm implemented in MATLAB’s Bioinformatics Toolbox. (55) To determine the sequence identity, alignment was performed using an identity mutation matrix, created using the eye(20) command. This amino acid mutation matrix used for global sequence alignment on the fasta sequences of peptides prior to TM-Score calculations. TM-Score matrix was computed for all the conformations (each peptide has five conformations). Conformations with structures greater than 0.5 are said to be within the same fold and thus similar. To select a non-redundant set of peptide sequences and folds we employed linear integer programming (ILP) based optimisation on a binary TM-score matrix weighted by the hotspot areas of all residues derived from their respective parent template (Σ_*r*_*P*_*r*_Δ*A*_*r*_). Dual-simplex algorithm with Gomory cuts was used to optimize the solution for a minimal peptide set:

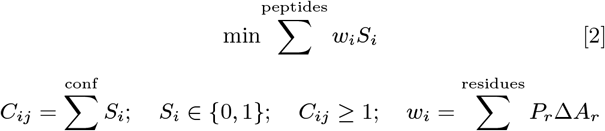

#### Peptide-NSP7 binding

All conformations in the non-redundant set were rigidly docked onto the binding partners of NSP12. The MM/PBSA-WSAS energies for binding were calculated for peptide-NSP12 complex. The binding similtitude of the peptides to partner PFIs was determined using a 70 length vector (an element for each residue) with respect to NSP7 by summing up the following for all docked poses of a given peptide-PFI pair:

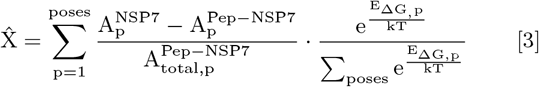

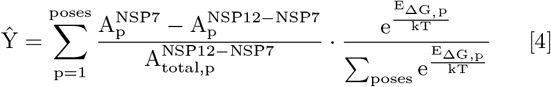

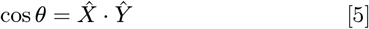

The cosine similarity of this vector with the vector obtained for each of the NSP12-NSP7 complexes (central structures using a backbone clustering of 2.5 Å) was found. Peptides with high dot products (> 0.90 of the maximum dot product) were docked using Rosetta’s FlexPepDock to introduce flexibility in the peptide backbones. The dot products were recalculated using the Rosetta interface energy scores (*I*_sc_) (56) to identify the most effective flexible peptides binding to NSP7. The top scoring structures from the enriched ensemble were subjected to explicit all-atom molecular dynamics simulations in the canoncial ensemble for 20 ns using the CHARMM-36m forcefield with the Nosé-Hoover thermostat and Parrinello-Rahman barostat.

### Experimental methods

#### SPR measurement

The binding Kinetics of NSP 7 with NSP 12, **P4** and **P5** proteins were measured by using a Biacore X-100 system with CM5 chips (Cytiva). The NSP 7 protein was immobilized on the chip by amine coupling with a concentration of 50 μg/ml (diluted by 0.1 mM NaAc, PH 4.5) according to the manufacturer’s recommendation. For all measurements, the same running buffer was used which consists of 20 mM HEPES, pH 7.5, 150 mM NaCl and 0.005% tween-20 with a flow rate of 30 mL/min at 25 degree C. Serially diluted protein samples are injected in a series of 0.19, 0.39, 0.78, 1.56 and 3.12μM with association time 60s and followed by 90s dissociation phase. The Multi-cycle binding kinetics was analyzed with the Biacore X-100 Evaluation Software (Cytiva) and fitted with a 1:1 binding model.

#### Competitive inhibition using ELISA

FLAG-taq Nsp12 and his-taq (HRP) nsp7 proteins of SARS-CoV-2 were purchased from BPS Biosciences, San Diego, CA, USA. The peptides were purchased from Genscript Biotech Corporation, New Jersey, USA. SARS-CoV-2 nsp12 protein was diluted at 10 ng/*μ*l-1 in PBS buffer. Two hundred nanogram protein was coated per well on a 96-microtiter ELISA plate (Nunc, Thermo Fisher Scientific) overnight at 4 °C. Next day, unbound protein was removed, and wells were washed thrice with 1X PBS buffer. Wells were then blocked with 4% (w/v) skimmed milk prepared in 1X PBS buffer and incubated at 37 °C for 45 min. The peptides were dissolved in water and were incubated with the coated protein of Nsp12 in an increasing gradient (1:1, 1:3, 1:5, 1:10, 1:20). For the positive control, no peptide was added to the well. Incubated peptides were allowed to interact with the coated NSp12 protein with slow shaking at room temperature for 1 h.Thirty microlitres of diluted nsp7 protein of SARS-CoV-2 (150 ng) was added to the well plate in triplicate and were allowed to interact with the coated Nsp12 protein at room temperature for 1 hr with shaking.Wells were then washed with 200 *μ*l of PBS buffer three times followed by incubation with 100 *μ*l of anti-his antibody prepared in 1X PBST buffer at 1:5,000 dilution and incubated at 37 °C for 30 min. The wells were then washed three times with 200 *μ*l of 1X PBS 2 buffer. One hundred microlitre 3,3’ ,5,5’ -tetramethylbenzidine substrate (Thermo Fisher Scientific) was added to each well and incubated for 10 min. The reaction was stopped by adding 100 *μ*L of 0.18 M sulphuric acid and the optical densities of the plate wells were measured using Biotek plate reader at 450 nm.

## Author Affiliations

Department of Chemical Engineering, Indian Institute of Technology Delhi, Delhi 110016, India

